# High Speed Visual Insect Swarm Tracker (Hi-VISTA) used to identify the effects of confinement on individual insect flight

**DOI:** 10.1101/2021.12.31.474665

**Authors:** Ishriak Ahmed, Imraan A. Faruque

**Affiliations:** Oklahoma State University, Stillwater, Oklahoma 74078

## Abstract

Individual insects flying in crowded assemblies perform complex aerial maneuvers by sensing and feeding back neighbor measurements to small changes in their wing motions. To understand the individual feedback rules that permit these fast, adaptive behaviors in group flight, a high-speed tracking system is needed capable of tracking both body motions and more subtle wing motion changes for multiple insects in simultaneous flight. This capability extends tracking beyond the previous focus on individual insects to multiple insects. This paper presents Hi-VISTA, which provides a capability to track wing and body motions of multiple insects using high speed cameras (9000 fps). Processing steps consist of automatic background identification, data association, hull reconstruction, segmentation, and feature measurement. To improve the biological relevance of laboratory experiments and develop a platform for interaction studies, this paper applies the Hi-VISTA measurement system to *Apis mellifera* foragers habituated to transit flights through a transparent tunnel. Binary statistical analysis (Welch’s t-test, Cohen’s d effect size) of 95 flight trajectories is presented, quantifying the differences between flights in an unobstructed tunnel and in a confined tunnel volume. The results indicate that body pitch angle, heading rate, flapping frequency, and vertical speed (heave) are all affected by confinement, and other flight variables show minor or statistically insignificant changes. These results form a baseline as swarm tracking and analysis begins to isolate the effects of neighbors from environment.

## 1. Introduction

The growing application of small-scale unmanned aerial systems has created a need for sensing and feedback paths that provide computationally-efficient, robust autonomy. The need for computationally-constrained robust autonomy is especially demanding in the case of small aerial platforms appropriate for swarm use, where size, weight, and power constraints limit the sensor payloads, processing, and communication tools that can be carried and near-neighbor interactions demand fast response times.

Insects are model systems for this challenge as they achieve robust maneuvers in unpredictable dynamic environment despite relatively limited neural resources. This performance includes multi-agent behaviors such as cohesion, swarming, and other coordinated motions involving navigation relative to each other. They often achieve these relative navigation tasks by means of implicit visual communication, i.e., without explicit communication links.

Despite these advantages, many attempts to replicate insect swarm behaviors have suffered from a lack of precise measurements quantifying their relative motion behaviors, and these bio-inspired routines are then inspired at the outline level rather than quantitatively consistent with experiments. The degree of biological consistency limits the resolution of the approaches; consequently, they often underperform the robustness seen in biological implementations.

Early work quantified the positions of insects as point masses, tracking only their positions. Recent high speed recording and visual tracking tools have enabled solitary insect measurements that include wing motions, which includes rigid and flexible body digitization of body and wing positions. Automated tracking has led to improvements in our understanding of the sensing and feedback paths used in individual flight control, including quantification of the flight stabilization reflex, and of the reward/penalty functions that insects feedback laws encode.

Detailed measurements of wing and body motion are needed in the multi-insect case to provide tools to extract the actual interaction rules implemented by swarming insects. These measurements need to precisely quantify the small perturbations in the moving motion, including increasing the tracked volume to allow for multiple interacting insects.

This paper introduces the first multi-insect, high speed tracker that simultaneously digitizes the flight trajectories of a flexible number of insects, including position and orientation for body and wings. High Speed Visual Insect Swarm Tracker (Hi-VISTA) is able to handle a flexible number and orientation of cameras and achieves improved throughput by parallel processing on several workstations.

In this paper Hi-VISTA is used to examine the effect of insect confinement. Although progress has been made in moving from tethered to free flight, the high lighting requirements and low depth of field of high speed cameras has typically limited measurement to small flight enclosures. The effect of these enclosures is not yet well quantified, but is needed to separate neighbour interactions from environmental responses.

The main contributions in this paper are a multi-insect tracker that provides measurements of body and wing states (position and orientation), and its application to a problems of contemporary relevance: the effect of confinement in flight. The study comprises 95 flight trajectories, with 15 variables tracked for each case, including body states and gross wing motion parameters such as stroke amplitude. The variables are analyzed by considering the trial wide mean and maximum values, and testing for statistical significance using Welch’s t-test (Welch, 1947), and Cohen’s d effect size.

## 2. Previous Work

In this section, previous insect trackers are reviewed, which are focused on either high speed tracking of an isolated insect including its wings, or lower speed tracking of the body states of multiple insects. Previous studies on enclosure and confinement are also reviewed for context of confinement experiment.

### 2.1. High Speed Wing and Body Solitary Tracking

In flight insect kinematics have been studied by resolving their body and wing orientation, and studying un-tethered flight normally uses a high speed multi camera setup to resolve wings and gain depth information. Early work used orthogonal cameras and manual methods to digitize several landmarks on a fly are digitized (Fry et al., 2003). The orthogonality restriction has been relaxed, largely by inclusion of a direct linear transformation (DLT) approach (Hedrick, 2008). Automatic trackers were then developed, including Hull Reconstruction Motion Tracking (HRMT) (Ristroph et al., 2009) which used orthogonal cameras to to produce maximally consistent 3D ‘volume pixels’ (voxels). Methodological improvements continued in Fontaine et al. (2009), which used model-based tracking and in Faruque and Humbert (2014); Kostreski (2012), which incorporated DLT coefficients to remove the orthogonality restriction and also included model based tracking with a Plucker line coordinate extended Kalman filter (EKF) formulation. Fontaine et al. (2009) manually initialized a predefined insect model and subsequently tracked. More recent work on larger insects has digitized the aerodynamic effects of deformable wing motions by including manually marked insects (Shumway et al., 2020).

### 2.2. Multi-agent body tracking

When multiple insects are present, previous work has focused on quantified body positions, using either an approach first tracks the insects in 2D camera views and then unifying the tracked trajectories, or an approach that reconstructs the 3D insect positions before tracking them.

3 camera views have been shown to be sufficient to simultaneously reconstruct large numbers of bat trajectories in offline processing (Wu et al., 2009). In this approach, 2D association relies on a consistency table and Kalman filter, and an iterative search procedure is then used to find a cost-minimizing correspondence between views. Similarly, large numbers (*>* 100) of flies have been tracked offline using 2 cameras by first finding the matching of targets in 2D, then finding the correspondence of the 2D paths and finally linking 3D segments by solving 3 linear assignment problems (Wu et al., 2011). Straw et al. (2011) implemented a real-time 3-D tracker (Flydra) for fruit flies and hummingbirds. This approach estimates the 3D position of the animal from multiple 2D camera views by the epipolar geometry intersection, nearest neighbor algorithm, and an extended Kalman filter (EKF). Grover et al. (2008) used multiple silhouettes to construct 3D visual hull of a fly and track the multiple hulls in real-time with a similar EKF implementation. Ardekani et al. (2013) developed a program to keep track of multiple flies in a 3D arena for long periods (hours), and classified behaviors via support vector machines.

Individual insects in these trackers can be identified after reconstruction using kinematic filtering techniques like the EKF, which has been employed in multiple investigations, or at the data-association level using color and form attributes (Kuo et al., 2010; Zhou et al., 2004). Body tracking of multiple flying animals has been shown to be a useful tool for identifying the interactions in aerial flight (Shelton et al., 2014).

In contrast to Section 2.1, these multi-insect tracker studies are limited to body position and orientation tracking without recorded information on wing motion. This approach leaves considerable ambiguity in which wing motions the animals use to effect those recorded body motions. The application to “black-box” modeling approaches at the individual feedback level requires detailed measurements of both inputs and outputs (Faruque et al., 2018). Consequently, the ability of these approaches to isolate the feedback control features, such as neighbor-modulated sensing and feedback gain adaptations, is limited.

We now discuss the relevance of the problems Hi-VISTA is applied to in this paper.

### 2.3. Confined flight

Insects reared and analyzed in a laboratory environment may not mimic naturalistic flight behaviors in free flight. The size of the laboratory flight enclosure used in insect flight experiments has long been known to affect the flight of the animal (Taylor, 1963). Stevenson et al. (1995) used cage size manipulation to demonstrate that the size of the enclosure affects hawkmoth forward flight speeds and results in wall-following behavior. It is likely that tethered flight induces some changes in wing kinematics, relative to naturalistic free flight behaviors and only in the last decade has detailed measurements of wing motions in un-tethered free flight become available (Ristroph et al., 2009; Kostreski, 2012; Faruque and Humbert, 2010).

While flight enclosure size and tethering may both induce artifacts in the resulting behavior, the effects of such laboratory preparations have received comparatively little study. In the last 30 years, tethered flight responses have dominated research. Various studies on flight mills over the last decade have been reviewed in Minter et al. (2018) and Naranjo (2019) regarding their methodology and approach. Flight mills have limits when it comes to analyzing outcomes and projecting them to the field because they are not natural (Minter et al., 2018). Because a tethered insect cannot carry its own body weight, abnormal flight behavior and erroneous reflections of natural flight performance may result (Dudley and Ellington, 1990). Several studies have indeed shown that laboratory insects show different behavior compared to wild ones (Baker et al., 1980; Nakamori and Simizu, 1983; Wales et al., 1985). Baker et al. (1981) reported that free-flying swarming locusts fly with higher wing beat frequencies and flight speeds than they do in laboratory experiments. Confinement size also affects the optic flow experienced by the insect, and it has been observed that insects such as bumblebees use different type of optic flow cues (lateral/ventral) depending on different tunnel sizes, and they prefer to fly over surfaces that provide stronger ventral optic flow (Linander et al., 2017). Insects such as orchid bees try to fly in a path which gives them maximal clearance from the edges which they determine relying on brightness cues (Baird and Dacke, 2016).

However, published literature lacks a direct comparison of the wing and body motions in free flight relative to those recorded in a confined enclosure. A major contribution of this paper is to use high speed tracking and statistical analysis to isolate the way in which these kinematics differ in response to changing enclosure size.

The remainder of this paper is structured as follows. Section 3 describes the construction of Hi-VISTA, including calibration, association, reconstruction, and segmentation of the reconstructed wings, body, and legs. Section 3.1.5 describes the method used to identify features on these bodies and measure their positions and orientations, and how it is validated with reference models in Section 3.6. Section 3.2 - 3.5 describes the experimental procedures and tools used. Section 4 report the tracker validation results and the main findings of the experiments.

## 3. Methods and approach

Hi-VISTA implements an association routine (Straw et al., 2011) and extends the capabilities of a single insect tracker to a multi-insect one to track the wing states as well (Faruque and Humbert, 2014). To avoid the necessity of an insect model, voxel based reconstruction methods (Kostreski, 2012) are incorporated because the multi insect environment holds a changing number of targets, including targets that are lost and reappear, often with differing body orientations. The implementation is focused on robustness to the variety of data measured in multi agent flight.

### 3.1. Multi insect tracker

This section describes the construction of Hi-VISTA. The main program is built in four major sections seen in Fig. 1 and elaborated in 3.1.2 - 3.1.5. A flowchart of the main program is shown in Fig. 1.

**Figure 1:**
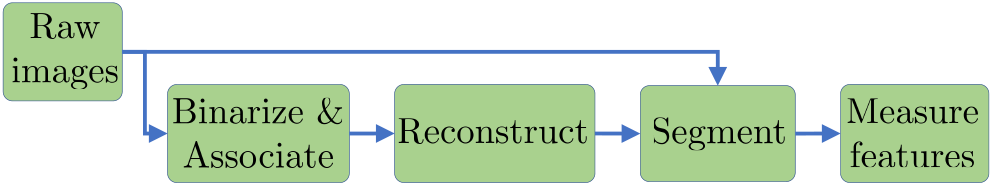
Flowchart of multi-insect tracking program

#### 3.3.1 Camera calibration

Camera calibration allows us to mathematically relate a 3D point in world to the 2D point in an image formed by the camera. The model used here is a pinhole camera, implemented through direct linear transformation (DLT) (Abdel-Aziz et al., 2015). The 3D *−*2D correspondence of a camera is modeled using a camera projection matrix *L* dependent on the camera optical properties, position, and orientation of the camera in the world coordinate system. The goal of multi-camera calibration program is to use simultaneous observations of the same point to estimate the individual DLT matrices *L*. As points in 3D world are quantized in pixels when the image is formed, *L* is not uniquely observable from a single point and so it is estimated by finding the optimum solution of a large dataset through bundle adjustment and random sampling consensus (RANSAC).

The camera projection equation here is

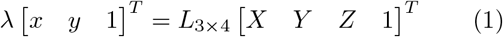

where, *L* is the DLT matrix, *x, y* are pixel co-ordinates of a point in 2-D image, *λ* is scaling factor, (*X, Y, Z*) is 3-D point in space.

The DLT matrices were estimated using a laser pointer to record individual points and RANdom Sampling Consensus (RANSAC) to identify the outliers and estimate the desired model, as implemented in Svoboda et al. (2005). This routine requires a single laser pointer to estimate the *L* matrices, which are then rotated to a desired world coordinate system as shown in Fig. 13.

#### 3.1.2. Binarize imagery & associate targets

This part of Hi-VISTA identifies one individual insect in different 2D camera views as shown in Fig. 2. Multiple insect centroids are identified in each view by subtracting the average background calculated from the images in each frame to detect the blobs.

**Figure 2:**
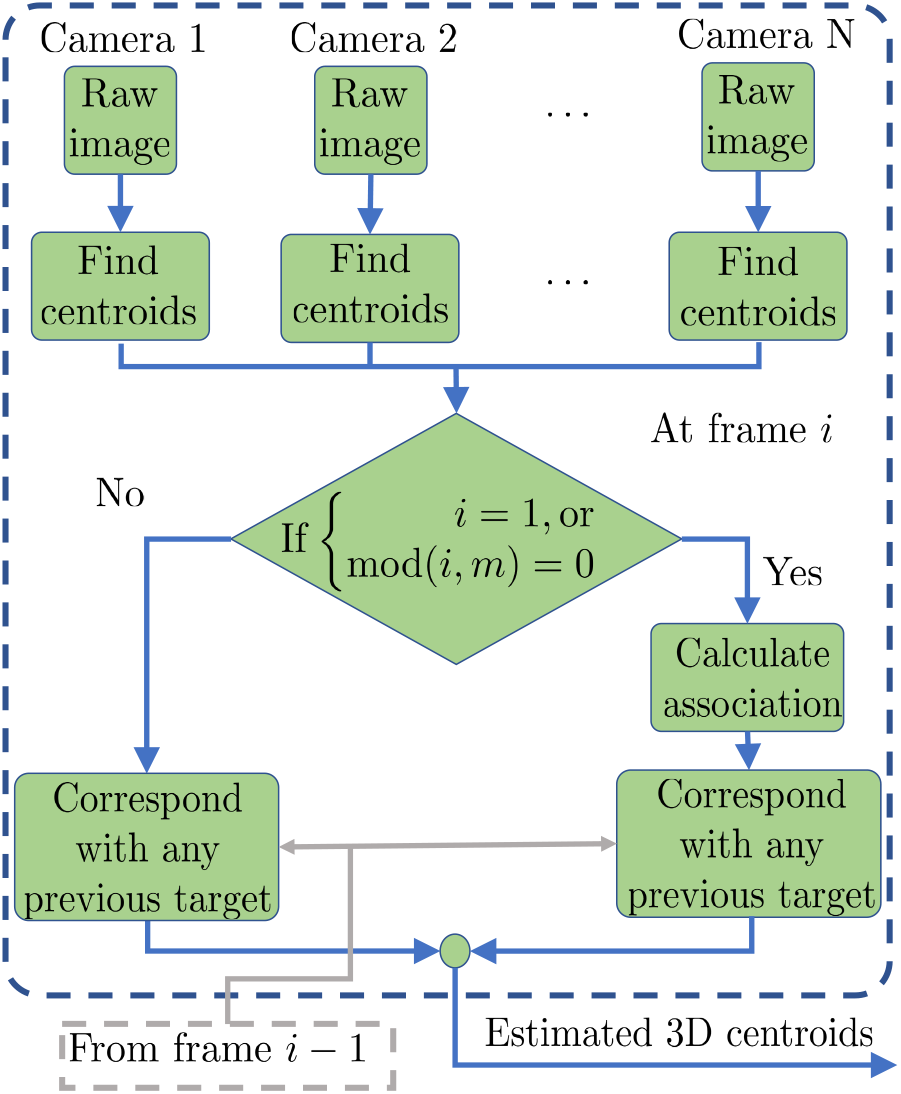
Insect association steps

Association is done by testing 3D points visible in the greatest number of cameras against a reprojection threshold as in the real-time (Straw et al., 2011), which is implemented in a MATLAB adaptation. This implementation is adapted for post-processing by including a capability to use different cameras for reconstruction and association tasks, subject to the limitation of visibility in more than 2 cameras. In this implementation assignment of multiple object ids to a single blob with nearest neighbour assignment corresponding to the case where the insect cover each other in a camera view was allowed which does not create a major problem if they can be separated in any other camera views because eventually the reconstruction routine can separate them in 3D.

For association every possible combination of triangulated 3D points from 2D centroids in multiple views is tested. Any 3D centroid visible in at least 2 cameras with a re-projection error below the desired resolution *η* is considered a valid association point.

In a *N* camera system a test combination of 2D centroids *C* is denoted by 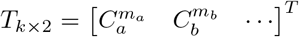 where, *a, b*, … are the camera indices and 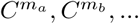 are one of the centroids visible in *a*^*th*^, *b*^*th*^, … camera and 2 ≤ *k ≤ N*. *T* is defined valid if every row *i* of the corresponding re-projection error vector *δ* has max_*i*=1,2,…, *k*_ *δ*_*i*_ *< η*, where,

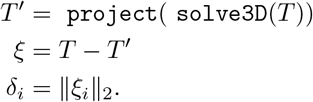

Here the function `project` uses camera projection equations (1) with 3D points to find the projected point in 2D, while `solve3D` uses camera projection with 2D points in available cameras to a solve for a least-squares-optimal 3D point.

With all valid combinations visible in the maximum number of cameras the association set *A* = *{T* ^1^, *T* ^2^, *… T*^*j*^*}* is computed where the frame contains maximum of *j* insects. The implementation supports arbitrary number of cameras with the flexibility to choose a subset of cameras if required for association and can keep track of insects visible in at least 2 cameras. After association matrix is computed, it is matched with any existing targets by minimizing target to target distance. Since *η* is a chosen variable, it is tuned specifically to the camera setup used.

#### 3.1.3. Insect Reconstruction

This part of the tracker generates the voxel reconstruction from the raw images and associated centroids as shown in Fig. 3,4. The observable 3D space can be discretized into volume pixels or voxels. With a valid associated centroid list available at each time step, every element in association set *A* is taken to generate estimated centroid *P* = solve3D(*T*). This point is then projected into voxel space, and a predefined scaler length, *d* obtained from the average size of a honey bee is used to define a search space such that *P* lies in the centroid of the cube with length *d*.

**Figure 3:**
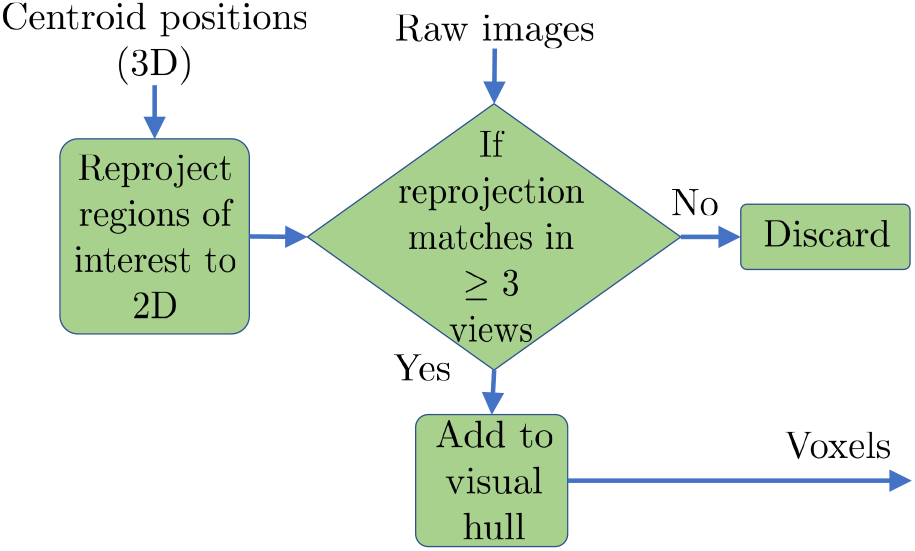
Insect reconstruction steps

**Figure 4:**
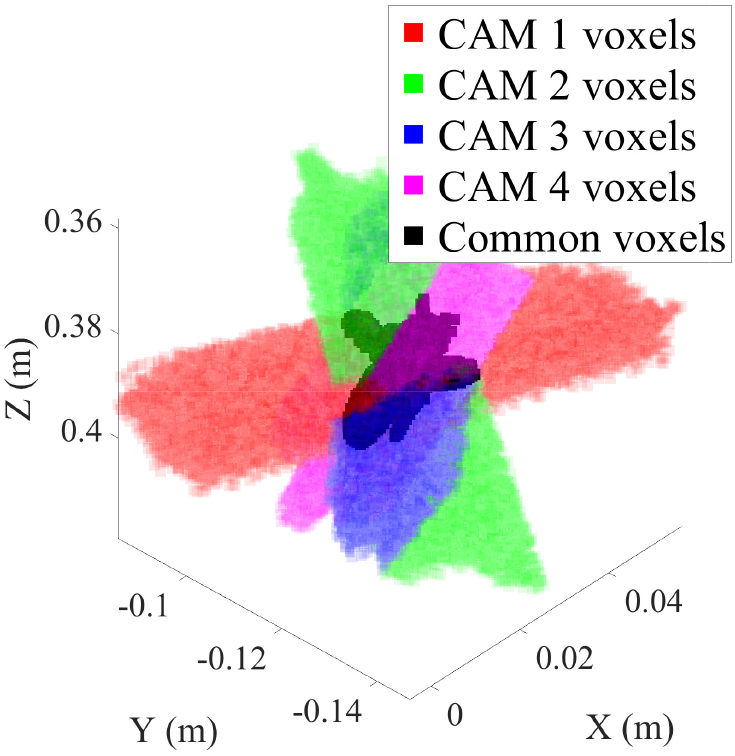
Insect is reconstructed by taking the common voxels in search area that could be projected back to insect pixels in the camera images

Every point in this search space that can be projected back on an insect blob in a preset number of camera views is registered as an insect voxel. This study’s implementation kept voxels which could be projected back on a binarized insect image in at least 3 views based on visibility in the cameras.

#### 3.1.4. Insect Segmentation

Segmentation refers to dividing the insect voxels into body and wing voxels as shown in Fig. 5. The intensities of identified insect pixels in each 2D view is used to do a preliminary segmentation that is refined in subsequent steps. In each view, the 2D insect pixels are clustered into two different groups where the darker and lighter pixels refer to the body and wing pixels. The histogram obtained from the pixel intensities as shown in Fig. 6 is used to estimate the probability density of intensity using a kernel density estimator (Hill, 1985). The probability density function generally shows two peaks and the value of normalized intensity at the minima between two peaks of the distribution is used as a segmentation threshold value to separate out wing pixel from body pixels. A voxel is assigned as a wing voxel if any of the views identify it as a wing, because the body is visible through the transparent wing. This purely intensity-based segmentation classifies voxels on the boundary of the body as wing voxels, and the segmentation must be refined by removing any wing pixel that is either close to the body and/or isolated. After distinguishing the wing/body voxels, *k*-means segmentation with *k*=2 is used to separate the two wings as in Fig. 7.

**Figure 5:**
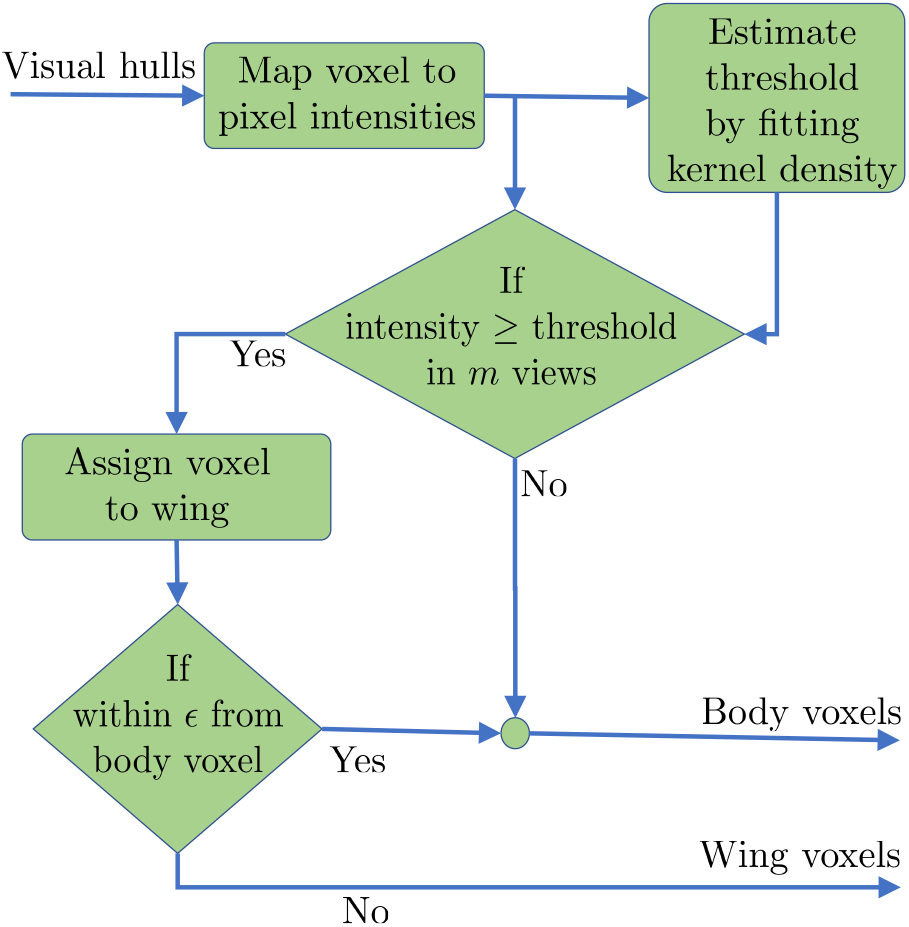
Segmentation to body/wings steps

**Figure 6:**
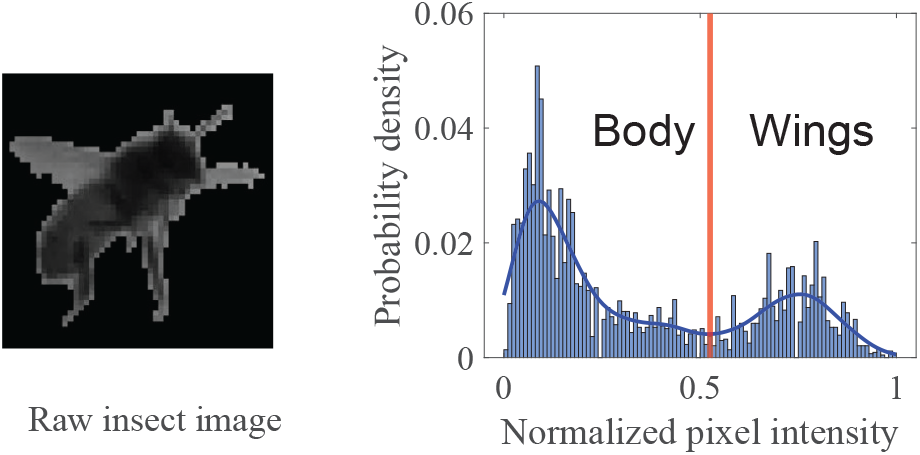
2D pixels on each insect body can be segmented to body and wing based on their intensity

**Figure 7:**
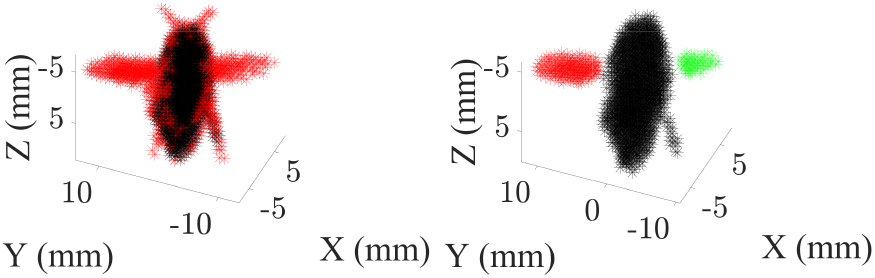
Purely intensity based segmentation (left) leaves ambiguous voxels close to the boundary of body voxels. Distance and connectivity measures are used to isolate wing and body voxels (right).

#### 3.1.5. Calculating insect parameters

Each insect is modeled as a 12-DOF system and the position of body center of mass, orientations of body and 2 wings are determined as shown in Fig. 10. Insect body parameters used in this paper are summarized in Table 1 and they are defined in Figs. 8 and 9.

**Table 1:**
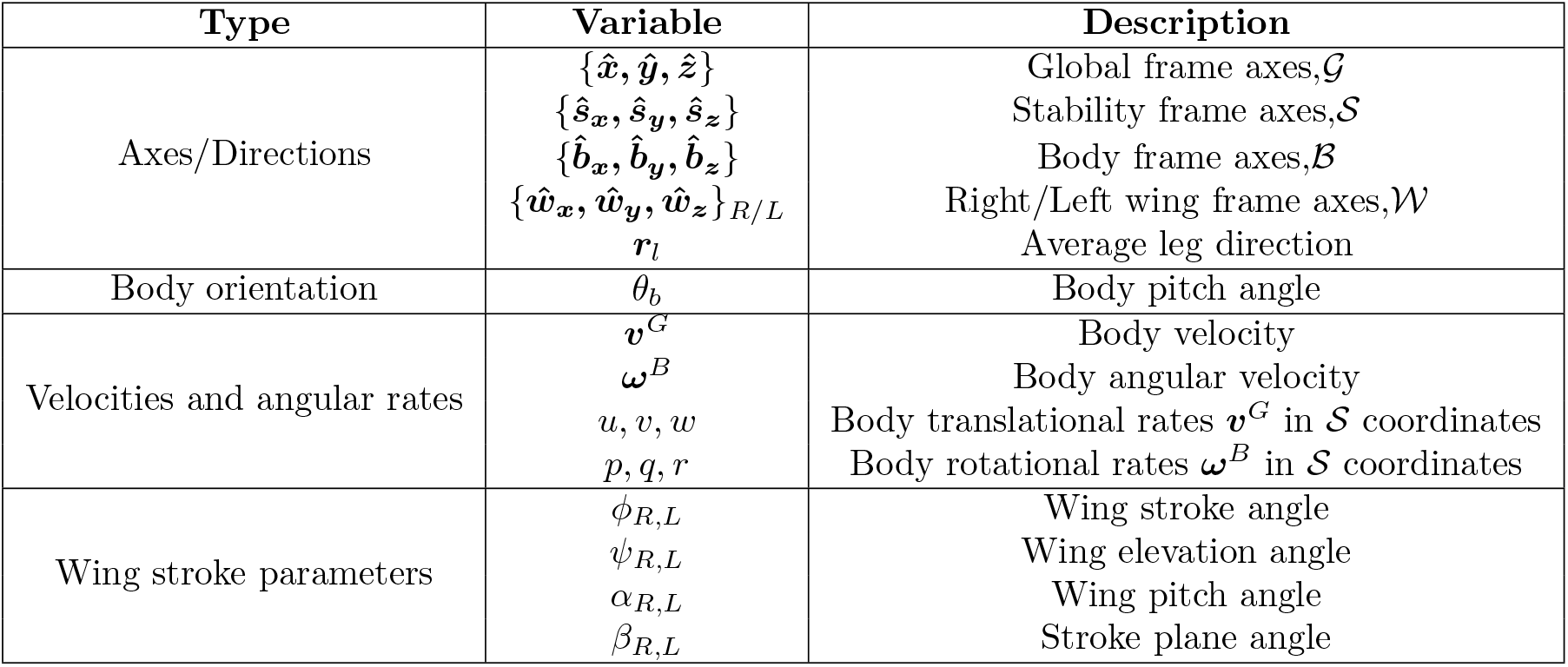
Notations used for defining insect pose

**Figure 8:**
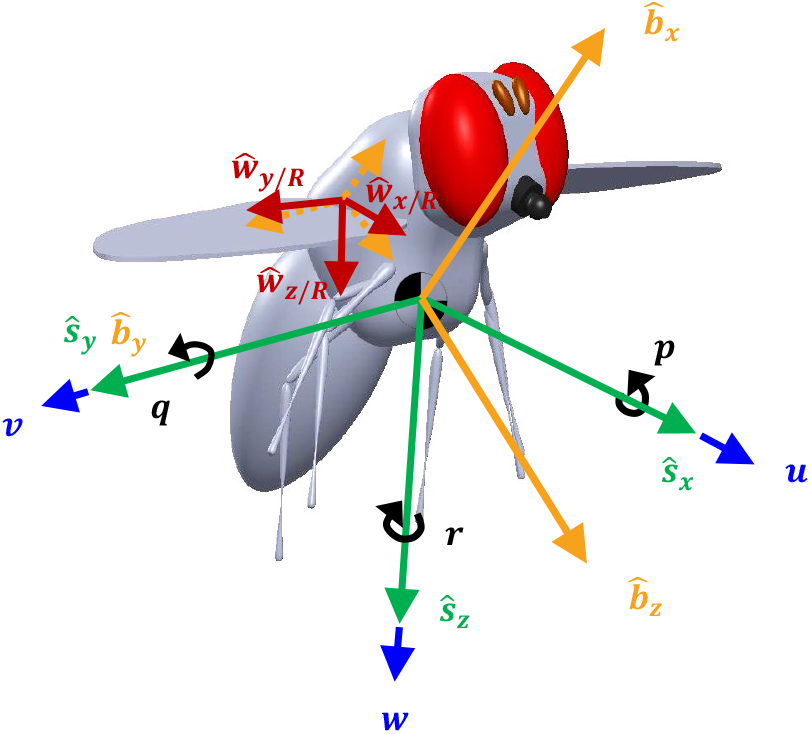
Body (orange) and stability (green) axes to define insect orientation.The right wing axes (red) are initially aligned to the body axes (dotted orange).

**Figure 9:**
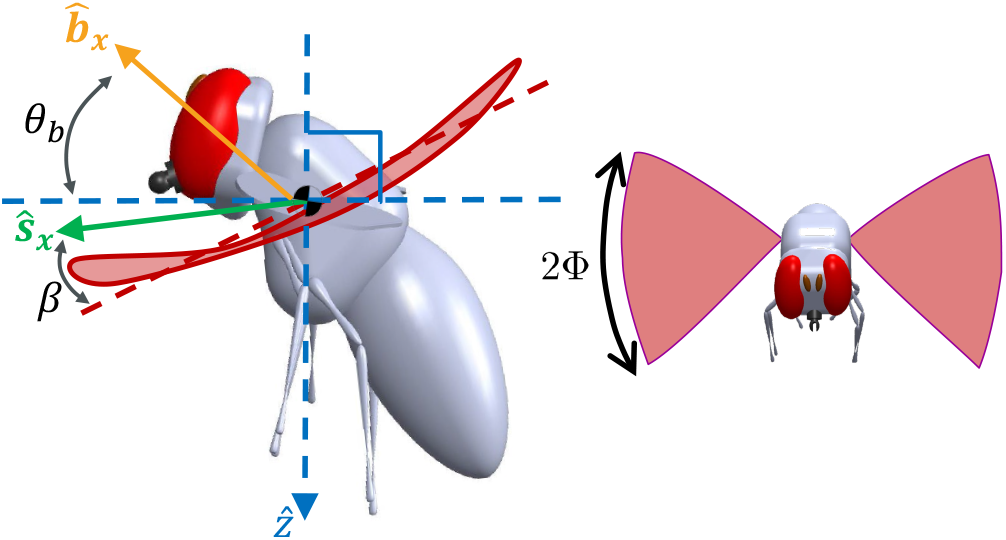
Wingbeat amplitude and body pitch angle definitions

**Figure 10:**
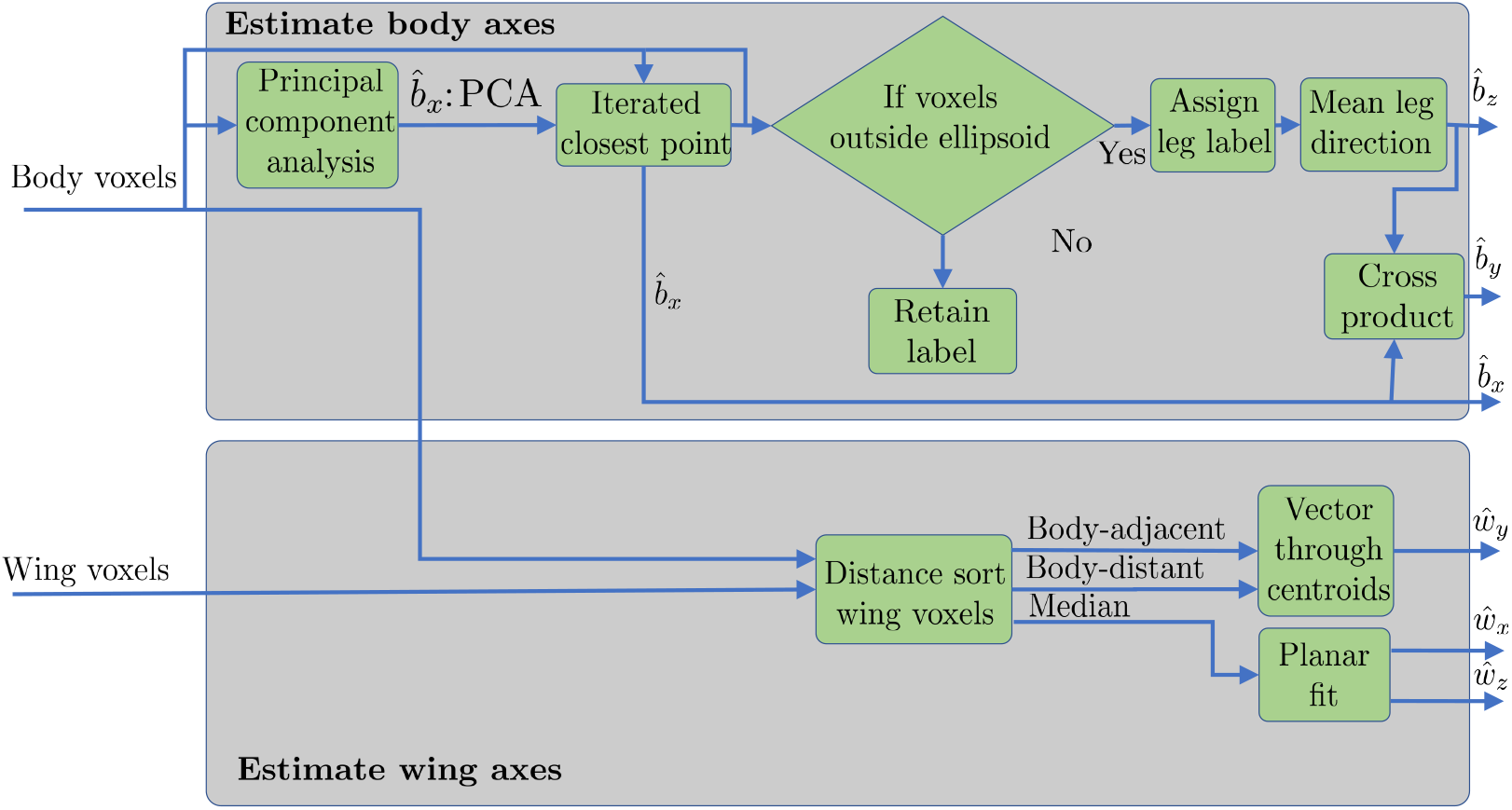
Insect parameters calculation steps

##### Identifying body frame

The body-fixed frame axes 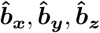 aligns with head-right-down, respectively. Insects’ generally prolate spheroidal bodies and axial symmetry about their body’s longitudinal axes results in the body roll angle being the most challenging feature to measure (Fontaine et al., 2009). Hi-VISTA uses visible legs as a reference to improve roll angle accuracy. The insect legs were considered to have free rotation about 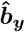. Leg direction was identified by an ellipsoidal fit with Iterated Closest Point (ICP) (Chen and Medioni, 1992) algorithm to the body voxel cloud, which computes a rotation matrix that minimizes the point to point distance in two point clouds. The ellipsoid major axis is initialized as collinear with the first principal component of body voxels. Voxels lying outside the ellipsoid are identified as leg voxels and the new ellipsoid major axis is considered as 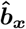 after the rotation. The centroid of leg voxels is then used to identify 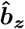 by reference to the vector ***r***_***l***_ from the body centroid to the leg centroid direction. 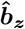 can be obtained by

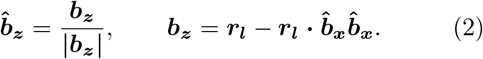

The body pitch angle *θ*_*b*_ is defined by the complementary angle of the angle between global vertical 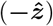 axis and body roll axis 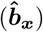 according to Fig. 9.

Hi-VISTA includes the capability to revert to previously methods of roll estimation that rely on assuming wing symmetry about the transverse body plane in cases of poor leg visibility (Fontaine et al., 2009).

##### Identifying wing frame

The wing frames are defined such that ***ŵ***_***y/R***,***L***_ aligns with the major wingspan. The nominal alignment as in Fig. 8 is chosen such that, right wing span vector 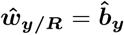 and normal vector 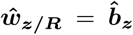 and for left wing left wing span vector 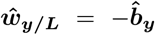 and normal vector 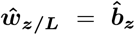. At each observation (time step), the wing span direction ***ŵ***_***y***_ is determined by taking the connecting vector between centroids of body adjacent and distant wing voxels. The wing normal vector ***ŵ***_***z***_ is estimated by the normal vector of the plane fitted through the remaining midspan wing voxels.

##### Determination of wingbeat frequency and amplitude

The wing angles are defined as the rotations needed to transform the body frame to a wing frame in each time instant, and described using a 3-1-2 Euler angle representation. The right wing angles *ϕ*_*R*_, *ψ*_*R*_, *α*_*R*_ were projected to planar motion (Faruque et al., 2018) using the stroke plane

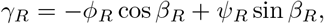

where *β*_*R*_ refers to the average stroke plane angle (Figure 9). The stroke plane is determined by linear fitting of *ϕ*_*R*_ and *ψ*_*R*_ over an stereotypical single wingbeat length. A fast Fourier transform of *γ*_*R*_ is then used to identify the wingstroke amplitude *Φ* and frequency *f* from the frequency domain peaks. Only the right wing is considered and it is assumed that left/right wing has same frequency (Deora et al., 2017).

### 3.2. Experimental setup and procedures

A T-shaped tunnel was built attached to a beehive of *Apis mellifera* residents and the two other legs exiting to the outdoors as seen in Fig. 12 and 13. Four Photron NOVA high speed cameras were used to film the intersection of the T-joint at 9000 Hz.

**Figure 11:**
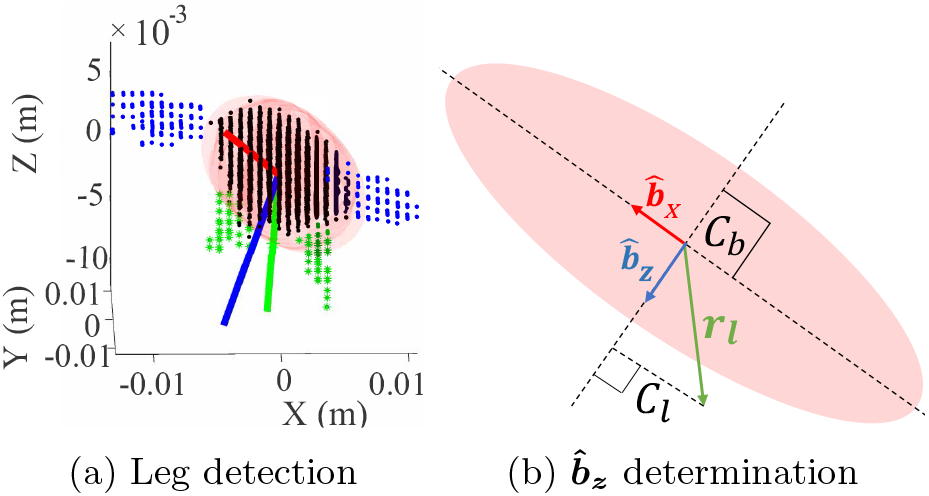
Reconstructed legs are used to identify roll angle through estimating 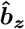. *C*_*b*_, *C*_*l*_ refers to body and leg centroids

**Figure 12:**
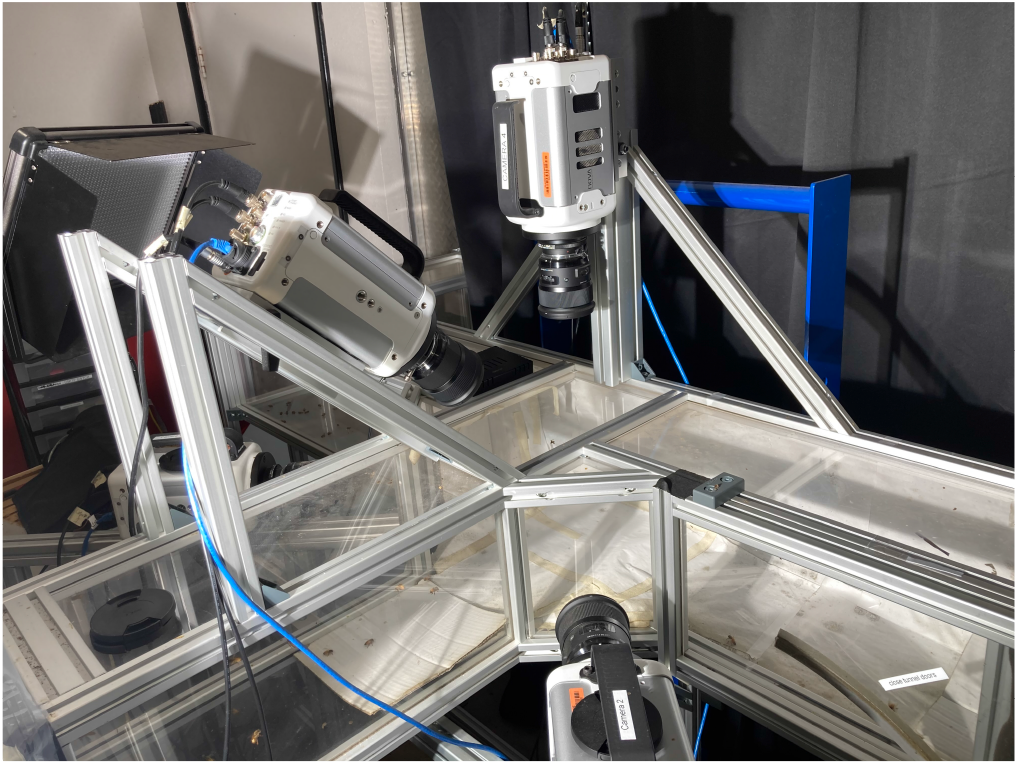
Tunnel setup attached to a beehive with camera setup to film the intersection

**Figure 13:**
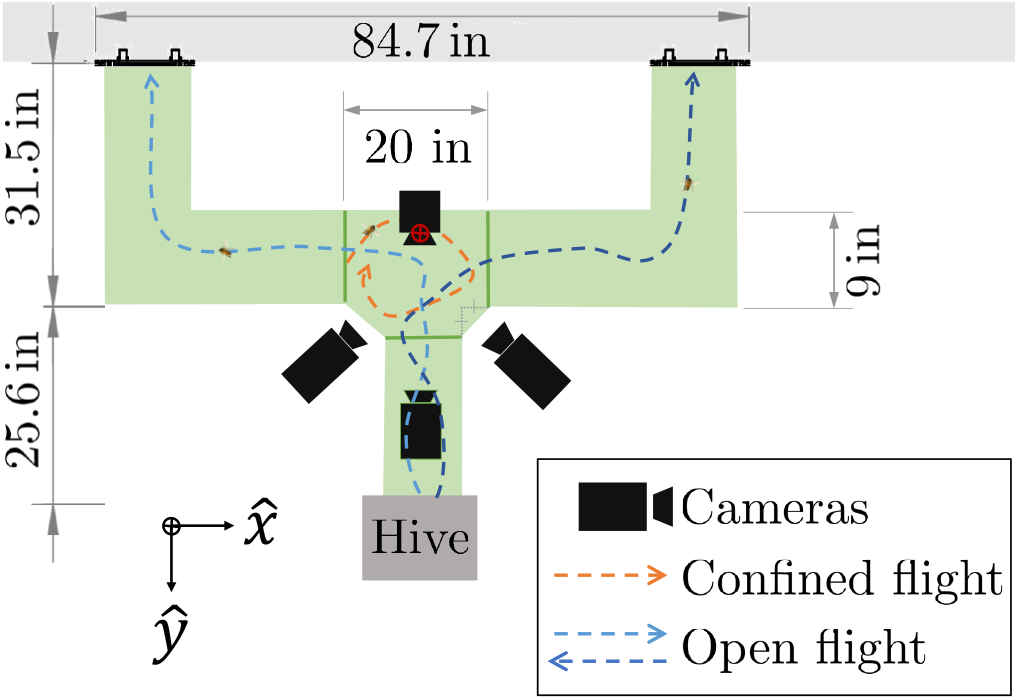
Insects fly through unobstructed tunnel in open tunnel experiments. In confined flights the intersection in confined by adding monochromatic partitions (showed in green).

### 3.3. Open and confined flights

Honey bee workers habituated to the experimental setup were filmed as they flew freely through the tunnel with no restrictions. The data was collected between 4 pm to 5 pm when the primary activity was foragers returning to the hive. In these trials multiple insects crossed the test section including a wide range of maneuvers. 60 flight sequences were analyzed for this study. Every captured video contained 2-4 insects in an observable volume of 875.67 *in*^3^. These flight trials are defined as “Open” for the rest of the paper.

“Confined” flight are the flight sequences taken while putting partitions in the intersection to trap insects. The enclosure volume was 2337.80 *in*^3^. 35 flight sequences were considered in this category. They were also filmed with the same camera covered volume of 875.67 *in*^3^.

### 3.4. Analysis procedure

For this study, the state variables in each flight sequence are represented by 15 (scalar) variables. For a recorded time history over [0, *T*_*r*_], where *T*_*r*_ is the time length recorded, time *t* was discretized as *t*_*i*_, *i* = 1, 2, 3…, *n* at a constant sample frequency, and the mean over a trial value of a measured variable *h*(*t*) measured the flight sequence was calculated as

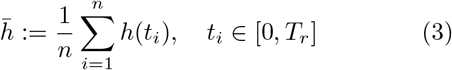

and the maximum value is defined as

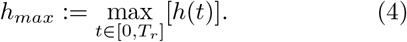

Each flight sequence was characterized by 15 scalar values as shown in Table 2. The set of these scalars is defined as *S*.

**Table 2:**
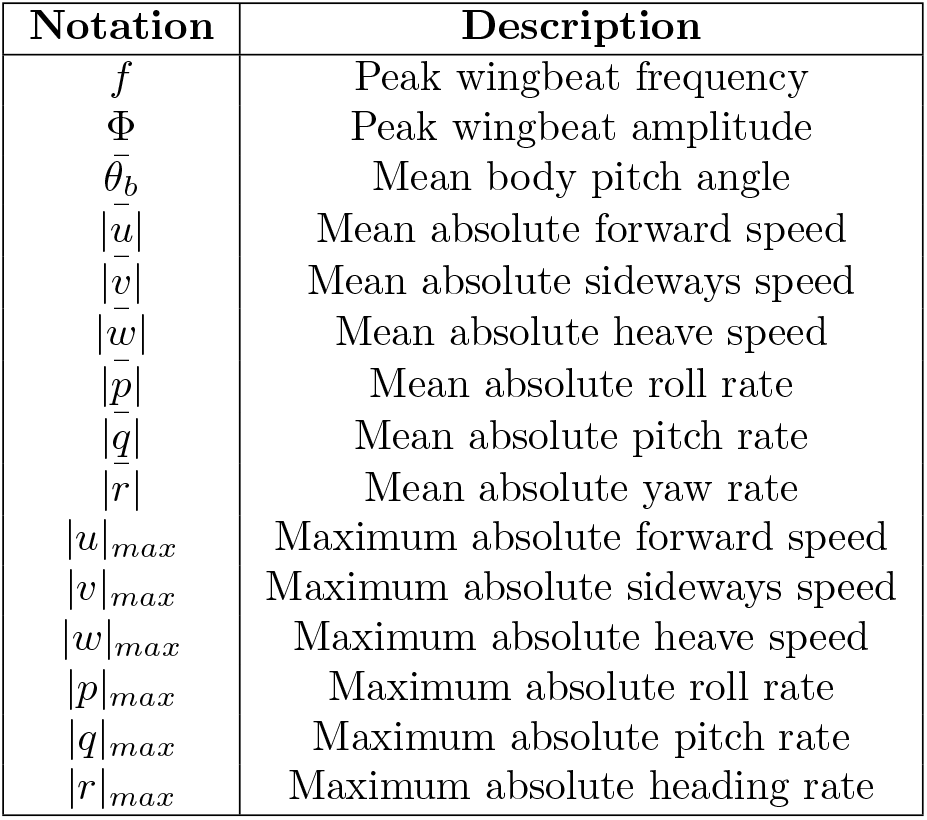
Characterizing variables in flight sequence

For each *s ∈ S*, population mean and standard deviation are considered for further analyses.

The population mean value of a variable was defined as

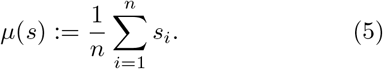

where *n* is the number of flight sequences recorded in the respective category (“Open” or “Confined”). The population standard deviation of a variable was defined as

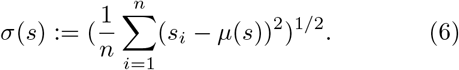

### 3.5. Statistical analysis tools

Binary statistical analysis was applied between the data of two groups “Open” and “Confined” to identify differences. The tools applied to this dataset were Cohen’s d effect size, and Welch’s t-test. Welch’s t-test tests the null hypothesis that two populations have equal means for some variable. This hypothesis was tested for each *s ∈ S* where the null hypothesis relates the open tunnel mean *µ*_*O*_ to the closed tunnel mean *µ*_*C*_ as

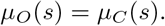

Student’s t-test assumes that the response variable residuals are normally distributed and the variances of populations are equal. Welch’s t-test does not assume equal variance and is helpful when the sample sizes are not equal. *p*-values are used to indicate the probability of the null-hypothesis being true. Cohen’s d quantifies effect size by indicating the shift of *µ*(*s*) in terms of pooled standard deviation. These tests are applied on the open and confined datasets having 60 and 35 sample points, respectively.

### 3.6. Validation

Camera calibration was validated by testing the individual reprojection errors. Insect reconstruction and feature estimation was validated by quantifying experimental error using at-scale fabricated reference models in known configurations as seen in Fig. 14.

**Figure 14:**
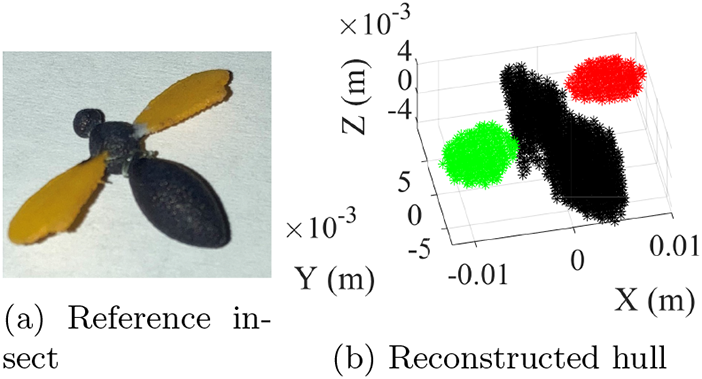
A reference insect (a) is reconstructed and the wing angles are compared for determining accuracy in the system. The insect and the reference insect are shown side by side for visual comparison

## 4. Results and Discussion

In this section, an example of tracked data and validation is presented, and each of the variables that show statistically significant deviations are examined.

### 4.1. Tracked data and validation

Camera calibration was accomplished providing camera projection matrices giving less than 0.5 pixels mean reprojection error over all views. An example of the tracked insect data output is shown in Fig. 15, which shows the planar motion of right wings for two digitized insects flying in close proximity to each other.

**Figure 15:**
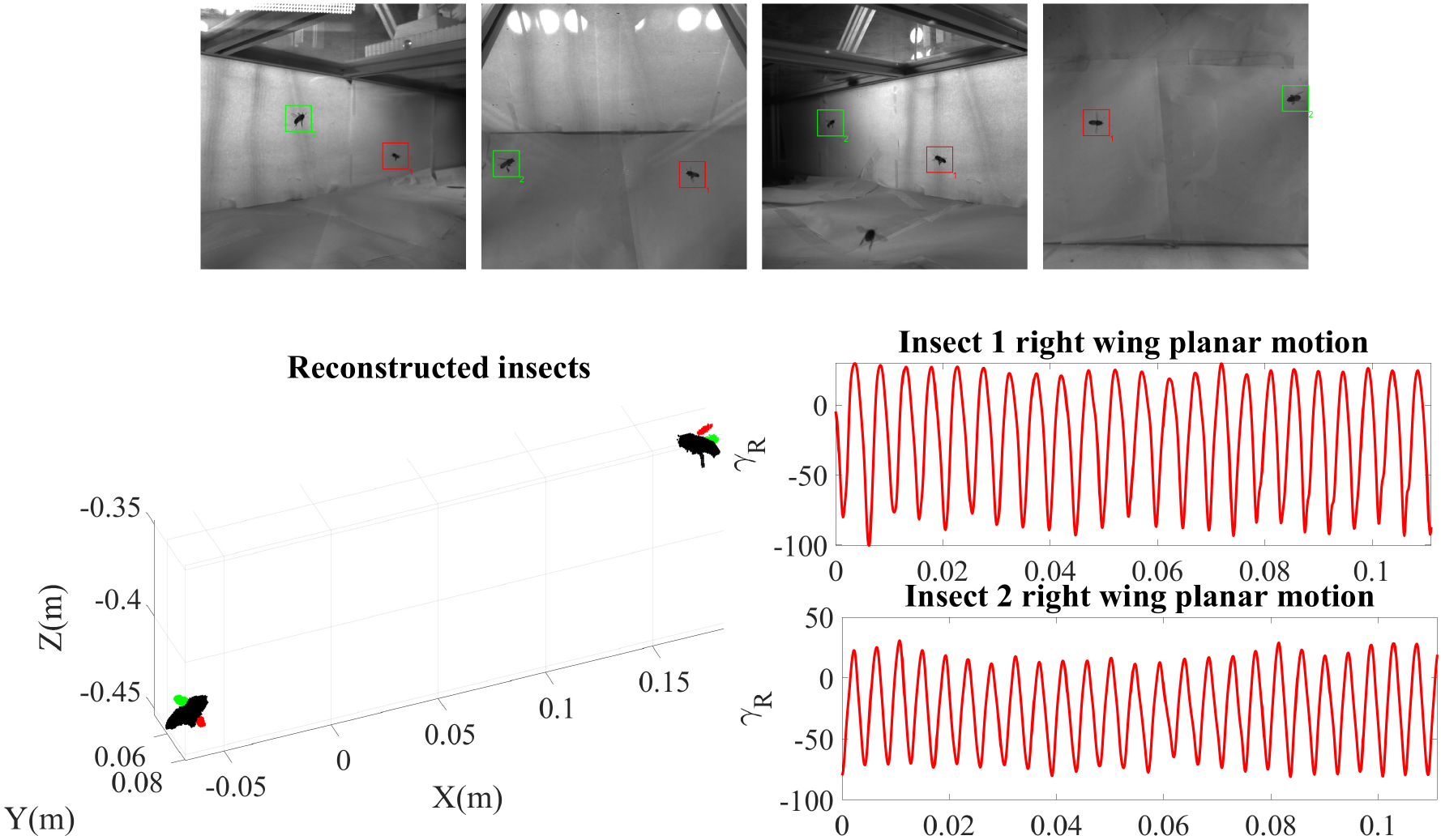
Example of output of Hi-VISTA

When the wing angles of a 3D printed insect were digitizing over 500 frames of motion, Hi-VISTA determined the 3-1-2 Euler angles as indicated in Table 3.

**Table 3:** Error in Euler (3,1,2) wing angle estimation for 3D printed insect

| Wing | Mean Error, ($^\circ$) | Std. Dev., ($^\circ$) |
| --- | --- | --- |
| Left | (-2.6, 0.6, 6.3) | (1.0, 1.3, 0.9) |
| Right | (-5.6, -3.4, 5.4) | (1.3, 0.8, 1.4) |

### 4.2. Difference between open and confined flight

Welch’s t-test applied to the p-values (Fig. 16) with *p <* 0.05 significance identifies 8 variables with significant differences in mean: 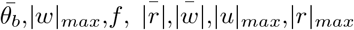 and 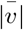. The Cohen’s *d* values quantifying variation size are shown in Fig. 17. 0.5 *<* |*d* |*<* 0.8 and |*d* |*≥*0.8 are used to identify as medium and large effects. A positive Cohen’s d value (*d >* 0) indicates confinement has increased the variable relative to “Open” flights.

**Figure 16:**
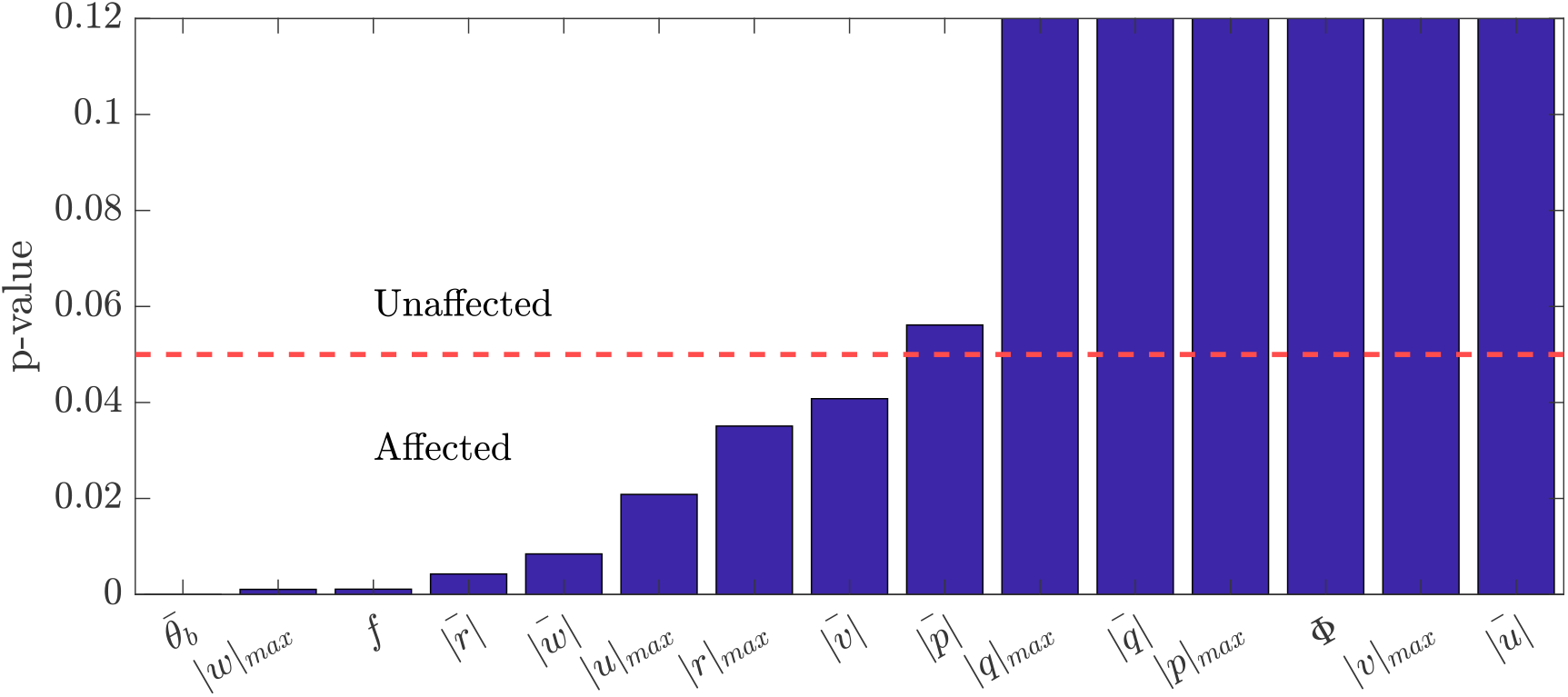
p-values (*<* 0.05) found with welch’s t-test determines the most affected variables

**Figure 17:**
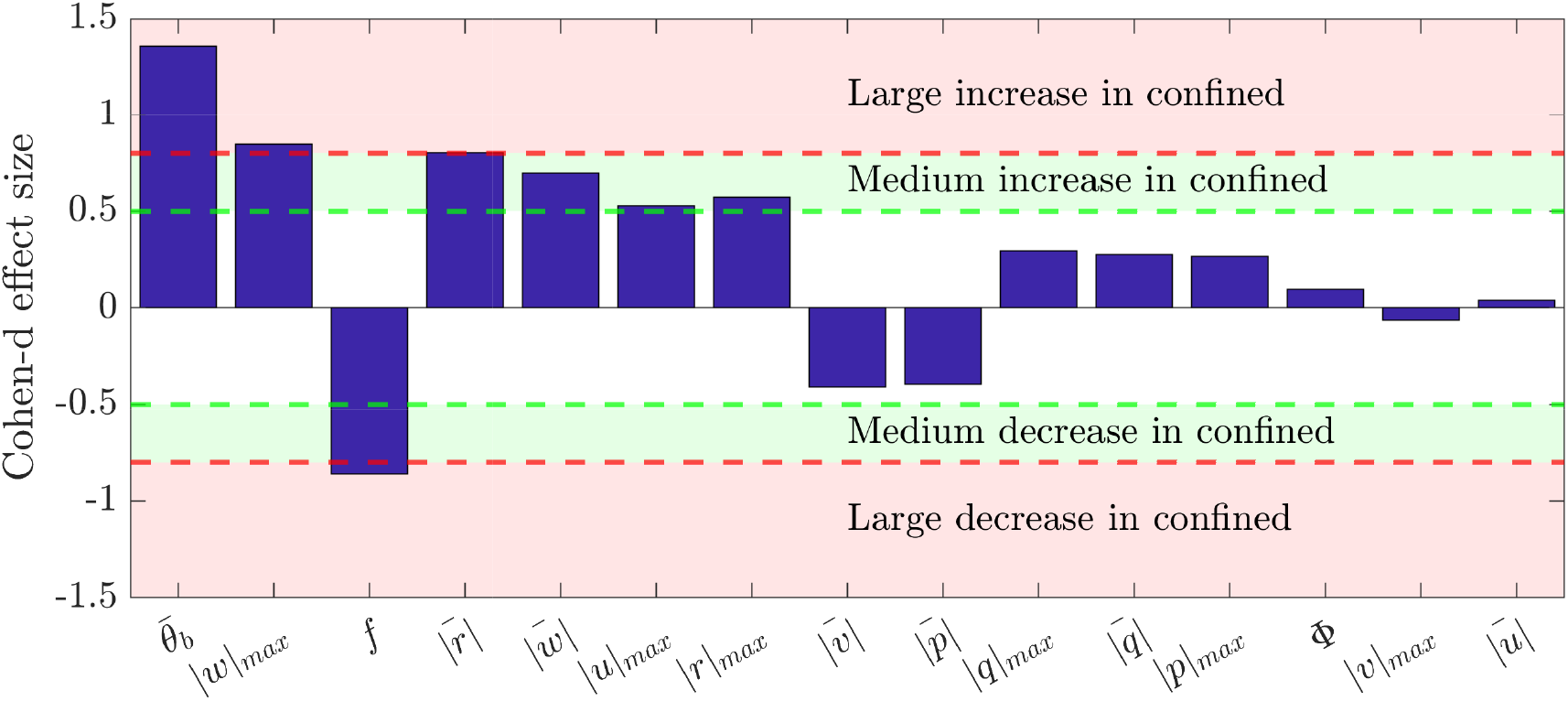
Cohen’s d effect sizes showing the relative shift of the mean of confined population with respect to the open population in terms of pooled standard deviations. 0.5 *<* |*d*| *<* 0.8 and |*d*|*≥* 0.8 are considered as medium and large effects.

#### 4.2.1. Body pitch angle 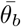

Body pitch angles during confined flight remained consistent with previous work of Vance et al. (2014). Mean body pitch angle *θ*_*b*_ significantly increases (*p <* 0.001, *d* = 1.3589) in confined flights relative to open flights. This increase may be partially explained by an increased occurrence of perching behaviors in confined flight, which often involve increasing pitch angle during the landing maneuver (Liu et al., 2019). In contrast, open tunnel flights contain a dominance of flights near cruise condition.

#### 4.2.2. Wingstroke frequency f

Welch’s *t*-test identifies wingstroke frequency as an affected variable (*p* = 0.0011, *d* = *−*0.8608). The mean frequency of wingbeat decreased significantly in the confined case. The frequency varied from 151 to 260 Hz in confinement versus 185 to 263 Hz in the open case. The asynchronous flight muscles in insects are generally tuned to operate near mechanical resonance (Josephson et al., 2000) and thus deviations from this frequency are often associated with reduced performance. Tethered flight has been associated with increase in wingstroke frequency and decrease in flight force output (Altshuler et al., 2005; Lehmann and Dickinson, 1997). In traditional insect flight enclosures, honeybees have been recorded in hover with relatively short amplitude and high frequency wingstrokes, using primarily amplitude tuning to generate their observed behavioral range (Altshuler et al., 2005; Vance et al., 2014).

#### 4.2.3. Heave velocity 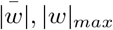

Confinement in-creased the variation in heave velocity (for 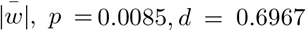 and for |*w*| _*max*_, *p* = 0.0011, *d* = 0.8475). Previous work on *Drosophila* showed that the insects track the altitude of a nearby horizontal edge (Straw et al., 2010). The confinement-related reduction in visual signals in this experiment may have inhibited the animal’s ability to fixate and track an edge, and thus impaired their ability to regulate altitude. For example, in contrast to Linander et al. (2017), no strong texture to provide ventral optic flow signals was provided.

#### 4.2.4. Heading rate 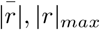

Confinement resulted in an increase in heading rate, both maximum (*p* = 0.0043, *d* = 0.8035) and mean (*p* = 0.0351, *d* = 0.5720) values. Previous work on tethered insects has indicated that saccadic turns induced during visual bar fixation have a lower magnitude than “spontaneous” saccades induced with a pseudo-random visual field (Mongeau and Frye, 2017). The monochromatic opaque enclosures used in confinement may reduce optic flow signals and inhibit visual fixation.

#### 4.2.5. Other variables affected

A weakly-associated increase (*p* = 0.0208, *d* = 0.5261) in maximum forward speed |*u* |_*max*_ was not analyzed further because the mean speed 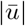 is not associated with a similar change (*p* = 0.8545, *d* = 0.0376).

### 4.3. Discussion of limitations & assumptions

The open tunnel tests are an improvement in biological relevance over contemporary laboratory experimental setups in literature quantifying insects in small clear enclosures, the current standard for “untethered free flight”. However, these insects are still restricted (mechanically, but not visually) to a tunnel environment and may show differences from flights in outdoor open spaces. In this study, manipulating both enclosure size and visibility, “open” tunnel must be interpreted relative to “confined,” and these results are relative differences that include both enclosure size and visibility effects.

If two insects are too near to each other such that the 2D blobs can not be distinguished in most camera views reconstruction may fail. This uncommon case is a fundamental limitation of camera-based measurement systems. The insect body reconstructor based on hull reconstruction is inherently limited in its ability to reconstruct micro details in a body. For example, the most prominent legs showed a strong influence over determining 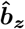, which is related to modeling the insect legs as having symmetry about the average leg direction. This approach performed well on the observed data set for a honey bee but may require adjustment for other species. The open tunnel experiments include flights of multiple subjects; this study’s statistical approach does not explicitly model interaction among subjects. Similarly, the binary analysis does not consider cross-corellations, and the effects may not be uncorrelated. Despite these limitations, the results serve the purpose of identifying the variables that must be interpreted with care to avoid conflating the effects of confinement and neighbor interaction.

### 4.4. Discussion

Detailed investigation of how insects modulate their sensing and feedback paths in response to nearby flying neighbors requires precise measurements of both wing and body motions for simultaneous insects. This paper developed Hi-VISTA, the first example of a high speed visual tracker capable of simultaneously measuring both wing and body motions for multiple insects in untethered flights. Hi-VISTA builds on existing work in visual hull reconstruction and target association to generalize contemporary approaches to existing work, and includes flexible camera number, orientation, and automatic background detection, including a new technique for measuring body roll angle. The performance measured on targets in known conditions is consistent with previous measurement systems (Kostreski, 2012; Ristroph et al., 2009; Faruque and Humbert, 2014).

Studies of insects interacting in confined spaces like laboratory test chambers require an understanding of how the behaviors are modulated by both confinement and the presence of neighbors, for example, metrics such as neighbor density. These factors have not been separated experimentally before, and in this paper, Hi-VISTA is applied to understanding the effects of enclosure and confinement.

Data points in ‘Open’ are taken from multiagent flights whereas the ‘confined’ data points are mostly from single insect flight. After partitions are applied after inserting insects in the filming volume, the tendency to fly simultaneously was low as per observation and it was, in a sense, helpful to draw a contrast between solitary confined flight and free group flight. The binary statistical analysis in this study identifies the subset of variables that could be targeted in future studies of varying enclosure size or insect density. The closed data is dominated by solitary flight, and can serve as a baseline for increasing numbers of agents in flight enclosure. The four variables identified (*θ*_*b*_, *r, w*, and *f*) are affected by confinement, and care must be taken not to attribute confinement-induced variation in these variables to the effects of neighbors in these studies. Conversely, variation in variables not affected in this study may be more likely to be associated with neighboring agents.

## 5. Conclusion

In this paper, a multi-insect tracker was developed which can be used to study both body and wing motion of insects enabling us to conduct more detailed studies on insect flight than the existing literature. This study is the first one comparing flight behaviors in confined insects and builds a foundation to more detail study on flight behavior. The insects flew at a higher pitch angle, with increased tendency of turning and moving in the vertical direction for which they had to use a lower wing frequency in the confined case. This preliminary study builds the groundwork for further involved studies with different types of insect flight and the multi-insect tracker gives an experimental environment to study multi-agent flight.

## Acknowledgment

This work was supported in part by ONR Young Investigator Award N00014-19-1-2216.

## Appendix A. Additional variables

The remainder of the variables analyzed in this study are included here, as quantified by their relative histograms.

**Figure A1:**
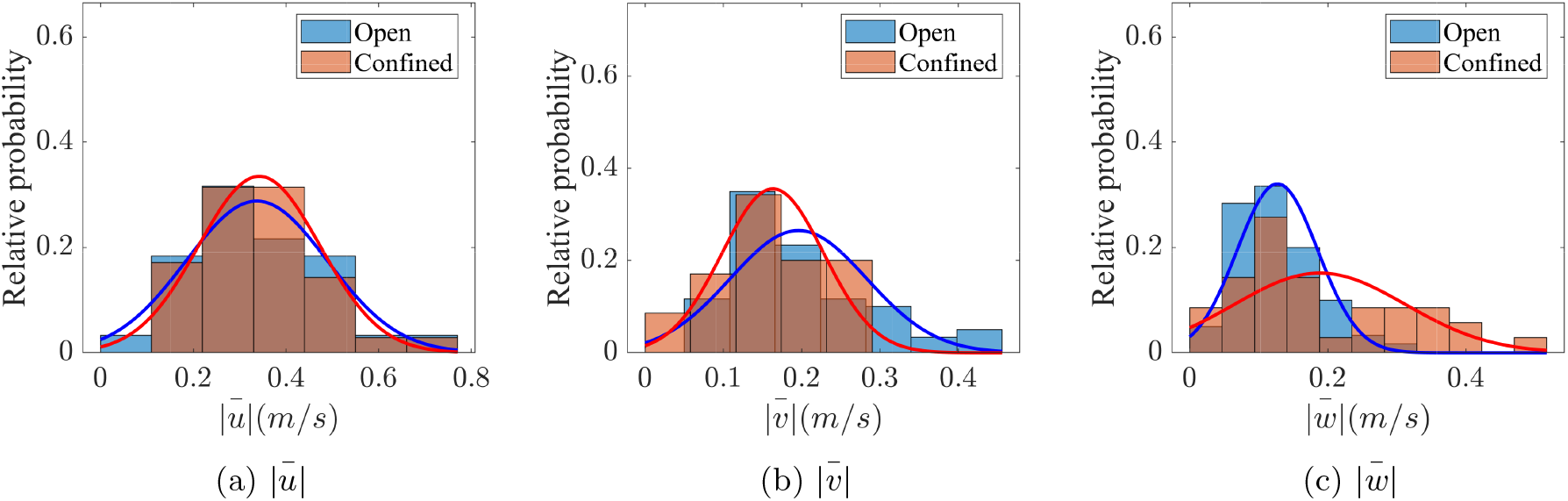
Mean absolute velocity histogram of open vs confined bees

**Figure A2:**
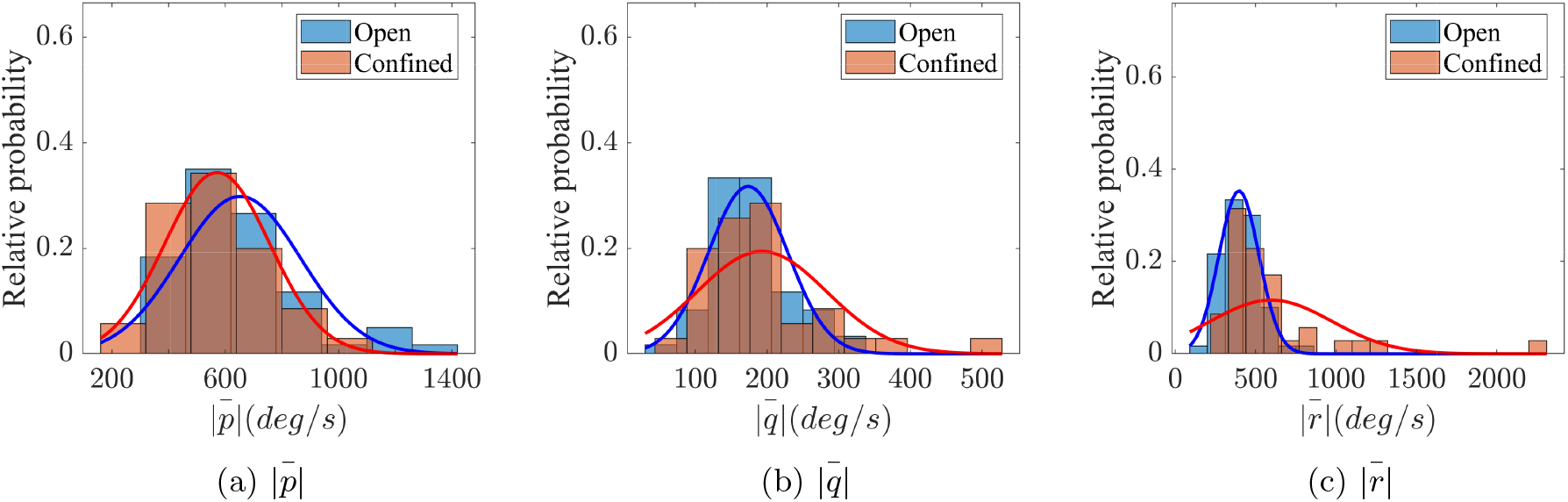
Mean absolute body rates histogram of open vs confined bees

**Figure A3:**
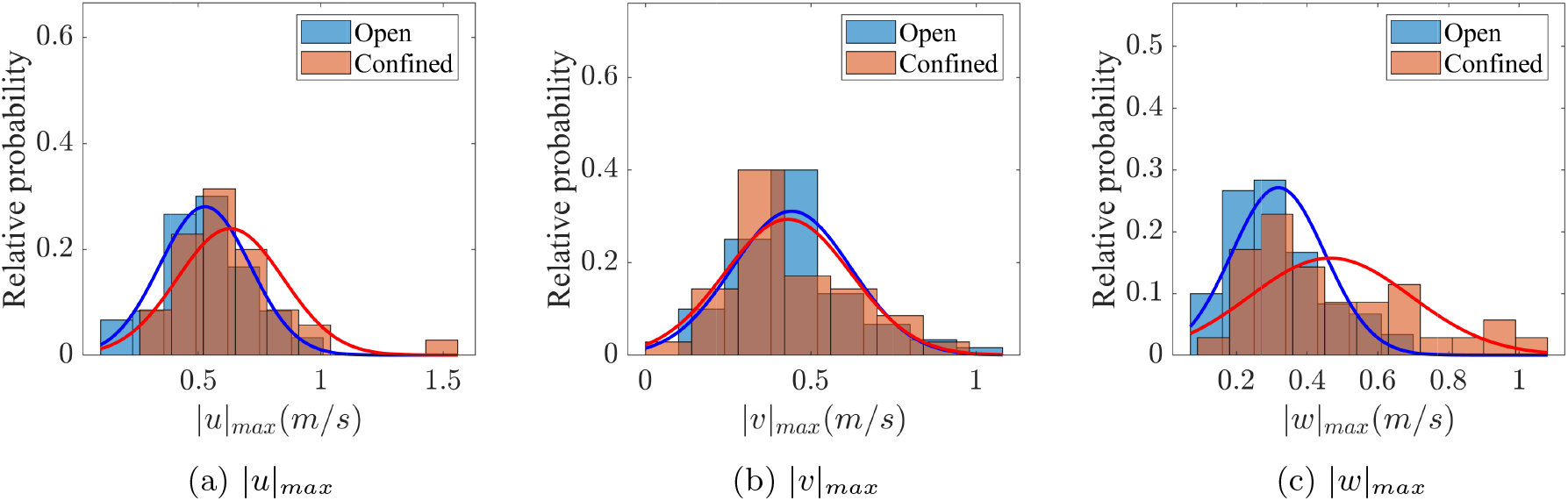
Max absolute velocity histogram of open vs confined bees

**Figure A4:**
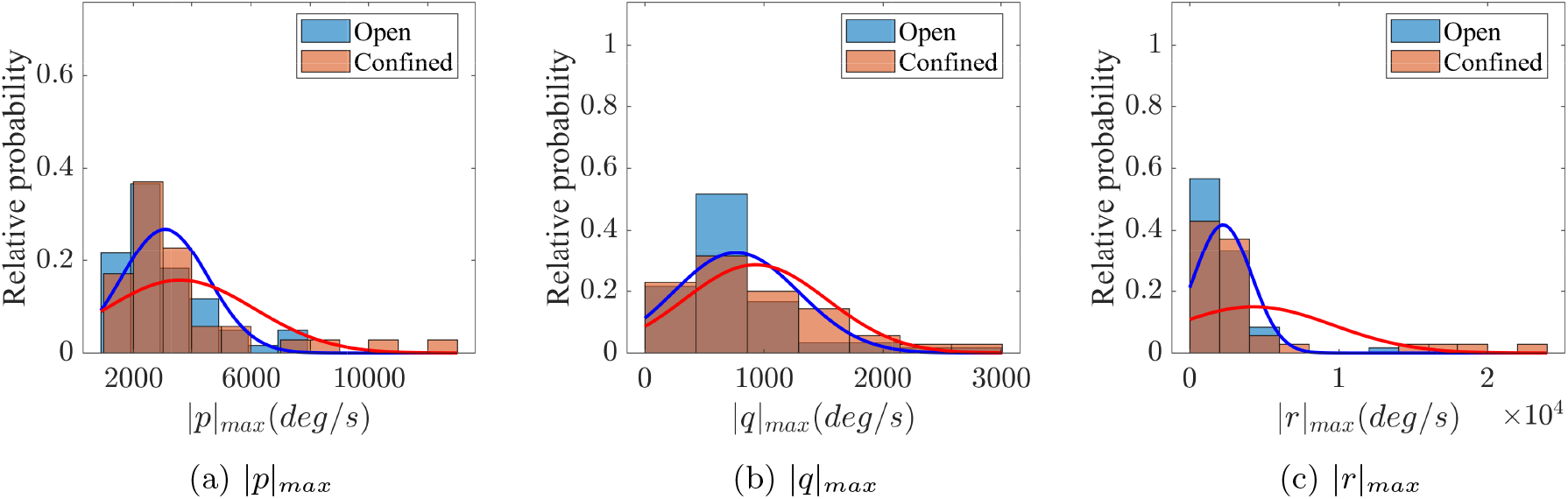
Max body rates histogram of open vs confined bees

**Figure A5:**
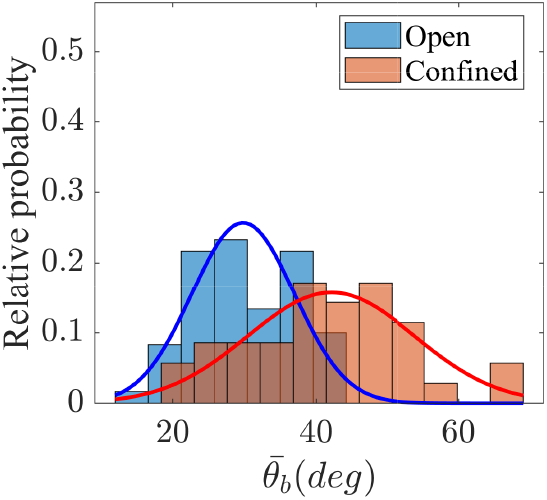
Body pitch rate histogram of open vs confined bees

**Figure A6:**
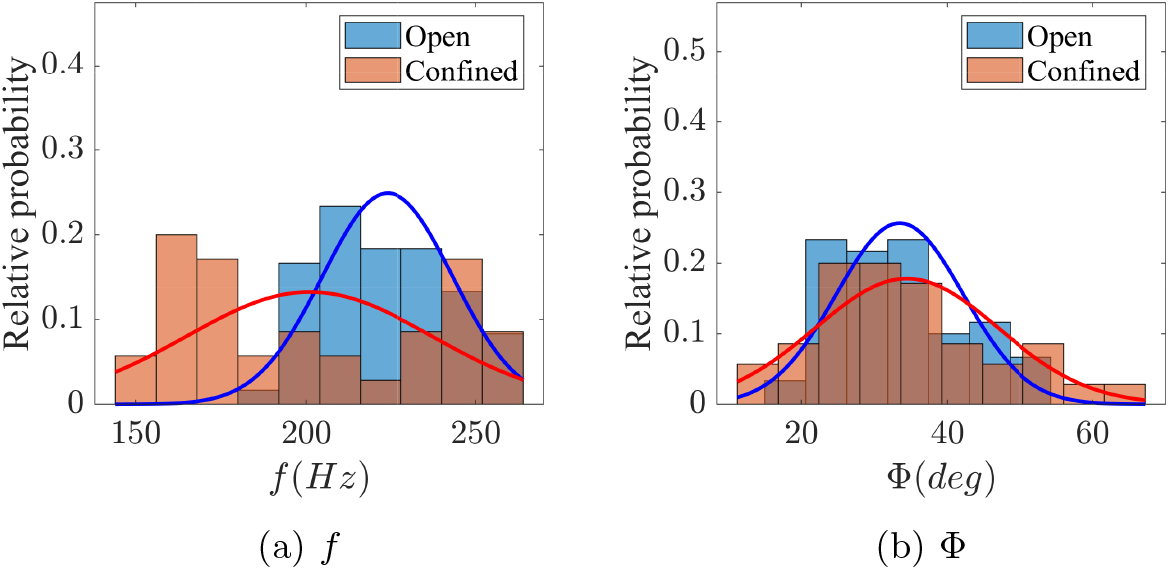
Wingbeat histogram of open vs confined bees

